# Parametric tests for Leave-One-Out Inter-Subject Correlations in fMRI provide adequate Type I error control while providing high sensitivity

**DOI:** 10.1101/2020.07.16.206235

**Authors:** L. De Angelis, V. Gazzola, C. Keysers

## Abstract

The inter-subject correlation (ISC) of fMRI data of different subjects performing the same task is a powerful way to localize and differentiate neural processes caused by a stimulus from those that spontaneously or idiosyncratically take place in each subject. The wider adoption of this method has however been impeded by the lack of widely available tools to assess the significance of the observed correlations. Several non-parametric approaches have been proposed, but these approaches are computationally intensive, challenging to implement, and sensitive methods to correct for multiple comparison across voxels are not yet well established. More widely available, and computationally simple, parametric methods have been criticized theoretically on the basis that dependencies in the data could inflate false positives. Here, we therefore endeavored to assess the actual performance of parametric tests on leave-one-out ISC values in two ways. First, we assess whether parametric tests protect against Type I error by assessing how often they find significant clusters of synchronized activity in publicly available datasets, in which such synchronization should not occur. This includes three resting state datasets, and one dataset in which participants did view movies, but where we randomly select segments that were not taken at the same time in the same movie. Contrary to what has been suspected, we find that parametric tests with corrections for multiple comparisons do protect appropriately against Type I error in that data. This was true for FDR correction at the voxel level at *q* < 0.05, with a minimum cluster-size of *k* = 20 voxels, FWE correction at the voxel level at *α* < 0.05, with a minimum cluster-size of *k* = 5, and for correction at the cluster-level with *p_unc_* < 0.001 with *k* = max(20, *FWEc*). Second, we assessed how these parametric tests compare with non-parametric methods when it comes to detecting ISC when participants actually did watch the same movies. We used a dataset including 150 participants viewing two movies, and used a bootstrapping thresholding of the ISC in the entire dataset to outline our best guess of the network of brain regions that truly synchronize while viewing the movies. We then drew subsamples of between 10 and 50 participants from the entire dataset, calculated the ISC, and thresholded it using our candidate methods. We find that FDR thresholding with *k* = 20 in particular, was substantially more sensitive than bootstrapping methods in detecting this network even in smallish samples of *N* = 20 participants typical of cognitive neuroscience studies, while at the same time retaining appropriate specificity. Because the parametric tests we show to perform well are more readily available to the neuroscience community than the non-parametric tests previously championed, we trust that this finding paves the way to a wider adoption of ISC, and empowers a wider range of neuroimagers to use ISC to tackle the challenges of naturalistic neuroscience. In particular in the context of often limited sample sizes and modest effect sizes in cognitive neuroscience, we trust that using FDR correction in particular will help neuroimagers identify the contribution of higher brain regions that process stimuli in more loosely timed fashions, more effectively than non-parametric alternatives.

## 1. Introduction

To overcome some of the limitations of traditional block- and event-related designs in functional magnetic resonance imaging (fMRI), the past two decades have seen the emergence of Inter-Subject Correlation (ISC) as an alternative method to localize brain regions involved in the processing of complex stimuli. This method leverages that voxels containing neurons with activity that is time-locked to a particular stimulus will show blood oxygenation level dependent (BOLD) signals that synchronize across multiple viewers of the same stimulus. Along the same logic, voxels that respond more reliably to one stimulus or condition than another will synchronize more to that stimulus or condition than to the other [1, 2, 3]. This method allows experimenters to use complex, long lasting stimuli, such as movies or stories, to probe the neural mechanisms underlying for instance emotional processing and social cognition [3, 4, 5, 1, 6, 7]. For traditional fMRI analyses, the field has converged onto highly validated statistical inference methods that correct for multiple comparisons across voxels in traditional contrast-based designs [8]. These “best-practice” methods are available in user-friendly software packages such as SPM and FSL [9, 10]. The adoption of these packages has tremendously improved the rigour and replicability of fMRI data-analyses by protecting neuroimagers from the dangers of writing their own analysis codes and of choosing sub-optimal methods. An issue hampering the wider adoption of ISC is that how best to determine whether a given pattern of correlations across participants and voxels is above chance or not remains a matter of debate. A successful approach to statistical inference in ISC has been to use pairwise correlations. Specifically, one correlates the brain activity of every possible pair of participants in each voxel, and then tries to infer whether these pair-wise correlations in one condition are larger than zero, or whether they are higher in one condition than in another. Because an experiment consisting of *N* participants generates a total of *N_p_* = *N*(*N* – 1)/2 possible combinations of pairs, with each participant’s data contributing to (*N* – 1) pairs, assuming that all of these pairs are statistically independent is untrue. As described in [11, 12], this is a case in which conventional parametric *t*-tests with DOF = *N*(*N* – 1)/2 – 1 do not appropriately control for Type I error, i.e., random data will generate significant *t*-tests at *p* < 0.05 in more than 5% of cases. For such pair-wise correlations, more appropriate non-parametric approaches should be pursued. Reasonably, since these non-parametric tests need to be performed in hundreds of thousands of voxels, corrections for multiple comparisons need to be applied [13]. Frameworks to apply such corrections in a way that takes the spatial auto-correlation of fMRI data into account however have not yet been optimized, and such tests are not available in the software packages in which most cognitive neuroscientists have been trained. Moreover, to perform such tests is computationally explosive because the number of possible pairs increases quadratically with sample size.

A second approach has been advocated, in which each participants time-course is not individually correlated with the time course of each other participant, but instead with the average time-course of all other participants (leave-one-out ISC). This approach considers the average brain activity of all other participants as a reference time-course that approximates the stimulus-triggered response in that voxel. The correlation of a participant’s time course with that reference time-course is then considered a measure of stimulus-triggered processing in that participant. This approach generates a single ISC map per participant, avoiding the artificial inflation of degrees of freedom encountered in pair-wise approaches, and leave-one-out ISC results have been analysed using traditional parametric *t*-tests after *z*–transforming the data by leveraging the power of standard fMRI analysis packages [3, 5, 14]. However, it has been argued, that such parametric approaches to leave-one-out data still run the risk of inflated false positives for two reasons. First, each subject does contribute to more than one ISC map, because it contributes to the average time course of all other participants, albeit in a way that is diluted by averaging [11]. Second, fMRI data follow a power law, and such data can generate spurious correlations [15]. However, whether these theoretical considerations actually translate into inflated Type I errors in ISC when applying parametric second level *t*-test to real fMRI data has never been explored. This is unfortunate, because using parametric *t*-tests would enable a wide community of neuroimagers to draw statistical inferences using software packages that have well validated corrections for multiple comparisons and lower the hurdle to the adoption of ISC analyses and the risk of errors associated with in-house analyses codes.

Here, we therefore assess the risk of false positive ISC results in parametric leave-one-out ISC analyses using publicly available resting state and task based fMRI datasets. For the resting state data, the rationale is that if we measure brain activity at rest, without participants viewing a common stimulus, we should not find significant ISC results, and all significant ISC can then be considered false positives. In addition, because the signal properties at rest may differ from those in the task-based fMRI applications in which ISC is typically used, we also evaluate the control of Type I error using a large dataset in which participants watched two movies. In this naturalistic viewing sample, we created cases where we do not expect significant ISC by calculating the ISC across different movies or different segments of the same movie. If parametric *t*-tests control Type I error despite the above-mentioned concerns, we would expect that when using corrections for multiple comparisons at *p* < 0.05 for family wise error corrections (FWE, be it at the voxel- or cluster-level) or *q* < 0.05 for false discovery rate corrections (FDR), a *t*-test on leave-one-out ISC on either dataset should not produce significant results in more than 5% of cases. We thus apply these analyses on 1000 subsamples of *N* participants from these larger datasets, and use SPM (www.fil.ion.ucl.ac.uk/spm), one of the most widely used and available fMRI analysis software to perform parametric *t*-tests on the ISC maps using the aforementioned corrections for multiple comparison and measure the false-positive rate for a number of combinations of *p/q*-thresholds and minimum cluster-sizes. From our data we conclude that both FDR with a minimum cluster size of *k* = 20 and FWE corrections with a minimum cluster size of *k* = 5 voxels provide robust protection against Type I error. Finally, we compared the sensitivity of these methods in detecting ISC in a dataset in which participants did watch the same movie, and synchronization is thus to be expected and compare these results with established non-parametric bootstraping methods [11]. We find that although all methods concord on the most highly correlated voxels, FDR is the most sensitive method, and is more sensitive that non-parametric bootstrapping despite its ability to control Type I error.

## 2. Methods

### 2.1. Datasets for false-positive analysis

To quantify the false-positives that arise when performing parametric tests on ISC data we analyze three publicly available datasets with resting state fMRI data as well as one fMRI dataset where participants engage in watching two different movies.

Two of the resting state datasets are from the Human Connectome Project (HCP) [16]: a first one consisting of 100 unrelated subjects (HCPS100) and a second one of 518 subjects including twins and non-twins siblings (HCPS500 release). The data from both HCP sets consists of 3T MR imaging data from healthy adult participants. The recordings include 1200 continuously acquired volumes for each participant, acquired using multiband acceleration at a TR = 0.72 s (multiband factor = 8) with a spatial resolution of 2 mm × 2 mm × 2 mm [16]. Additional behavioral and demographic measures on the individual participants can be downloaded from the project website [16].

We include a third resting state dataset from the Autism Brain Imaging Data Exchange (ABIDE) [17]. From this dataset we retrieved 100 recordings from healthy participants used as control group for the ABIDE study. The data was collected for ABIDE by the New York University Langone Medical Center. This dataset includes a minimum of 176 continuously acquired volumes for each participant acquired at TR = 2 s with a spatial resolution of 3 mm × 3 mm × 3 mm [18]. The inclusion of datasets aquired at different TR values (0.72 s for HCP and 2 s for ABIDE) ensures that our findings apply to a broader range of datasets.

The fourth dataset consists of two naturalistic viewing fMRI scans from the Healty Brain Network (HBN) [19]. From this dataset we selected 3T fMRI recordings from 150 healty subjects watching the movies “Despicable Me” (DM) and “The Present” (TP). We selected the 150 subjects by filtering the recordings from the sites CBIC and RU corresponding to participants older than 15 years. In this way we ensured a reasonable quality for the registration to a common atlas. For each participant we have two fMRI recordings, including 750 continuously acquired volumes for DM and 250 for TP. For both movies, the data has a spatial resolution of 2.4 mm × 2.4 mm × 2.4 mm and a repetition time TR = 0.8 s (32-channel head coil and CMRR simultaneous multi-slice echo planar imaging sequence). Additional imaging and phenotypic data on the individual participants can be downloaded from the project website [19].

For the HCPS100, HCPS500 and the ABIDE data, we downloaded and used the preprocessed datasets. Details on the specific preprocessing can thus be found in [18, 20]. We did not apply additional processing prior to the inter-subject correlation analysis. Briefly, the preprocessing that had been applied to these datasets involved standard pipelines combining FSL [21], FreeSurfer [22], Connectome Workbench [23], AFNI [24] and ANTS [25]. The imaging data from the ABIDE dataset had been registered to a MNI template with isotropic voxel size of 3 mm and spatially smoothed with a 6 mm FWHM Gaussian kernel. The HCP dataset had been registered on a MNI template with isotropic voxel size of 2 mm without smoothing. It should be noted, that the HPC500 dataset contains siblings that could have brain activity that is more similar than that of non-siblings. We however ignored this fact in our analyses.

For the HBN dataset the data needed preprocessing. In particular, for performing ISC analysis it is essential that each recording is registered to a common atlas. In order to keep every step performed in our study as standard as possible, we performed preprocessing with *fmriprep*, a robust and automatized preprocessing pipeline [26]. With fmriprep, we registered each functional recording to a MNI template, with isotropic voxel size of 2.4 mm. Additionally, we spatially smoothed the data with a 6 mm FWHM Gaussian kernel

### 2.2. False-positive analysis strategy

For each of these four sets, we perform 1000 analyses by randomly subsampling a certain number of subjects. We subsample groups of *N* = 10, 20, 30, 40, 50 subjects, with the aim of emulating realistic numbers of participants in an fMRI experiment. Having defined a sample of *N* subjects, we extract a time series from each participant (Fig. 1). The HCP dataset consists of functional files of approximately 1 Gb per subject containing 1200 volumes. To limit the memory resources required to carry our false positive analyses, we extracted 200 contiguous volumes and perform leave-one-out ISC across these 4D volumes. From the 1200 volumes available for each subject we extract the 200 in the middle (volumes [500:699]). To ascertain that our findings are not restricted to this arbitrary choice of 200 volumes, we also replicated our analyses for the sample size most typical of contemporary fMRI studies (*N* = 20) using the following numbers of consecutive volumes: 10, 20, 50, 100, 200 and 400 (see Supplementary Material 1). This analysis came to the same conclusions as those presented in the main paper. For the ABIDE dataset we used the full 4D volume, i.e., volumes [0:175] from the 176 volumes available for each participant.

**Figure 1:**
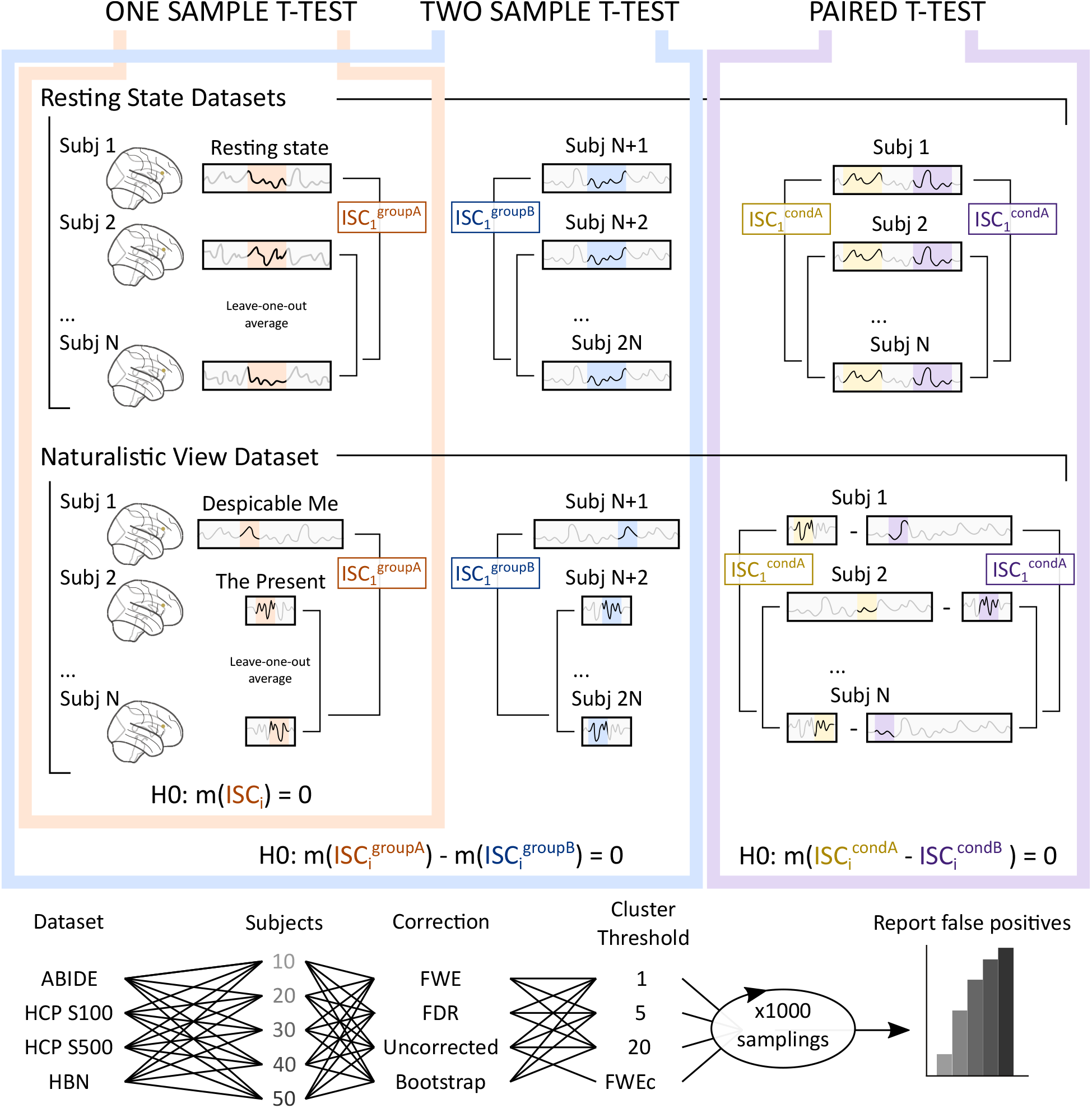
Analytical procedure for false positive rate quantification: Calculating leave-one-out ISC value involves (i) selecting a subsample of *N* subjects from the entire dataset, (ii) taking one subject out, (iii) averaging the time course of the other subjects, and (iv) calculating the Pearson *ρ* value between the left-out participant and the average of the others. For the one sample *t*-test (left), for the resting state datasets (top) this can be applied directly using a fixed section of each participant’s resting state time-course (marked in skin colour), because these time-courses should not be synchronized. The resulting ISC values across the *N* participants and many voxels are compared against *H*_0_ : *m*(ISC_*i*_) = 0 using a parametric *t*–test. For the HBN dataset, the actual time courses are synchronized in some voxels because participants were watching the same movie and significant ISC would thus not be false-positives. To assess false-positives, we disturb this temporal association while preserving the other properties of task-based fMRI signals. Accordingly, for this dataset (middle row), we select a random epoch from one movie (DM) for the left-out participant, and select random sections of the other movie (TP) to average across the other participants. This ensures that the selected segments should no longer be systematically synchronized, and significant detected ISC can be considered false positives. For the two sample *t*-test (center panel), in addition to what is already done for the one sample *t*–test, the same is performed in another subsample of *N* non-overlapping participants. This generates ISC values for two groups (A in skin colour, and B in blue), and the resulting ISC values are contrasted against the two groups in a two sample *t*-test. Finally, for the paired t-test, we again use two different procedures for the resting state and naturalistic view datasets. For the resting state dataset, we select two non-overlapping segments in the resting state time-series of each participant. The first (yellow) is analysed across participants to generate ISC values for each subject *i* in condition A. The second (lavender) is analysed across participants to generate ISC values for each subject *i* in condition B. The two ISC values of each participant are then subtracted, and these differences tested against 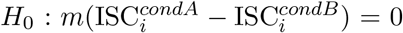. For the naturalistic view dataset, we randomly selected a segment in movie DM and one in movie TP for each participant. We then randomly attribute one of the two to condition A (yellow) and one to condition B (lavender), separately for each participant. We then calculate the ISC across all the condition A, and do the same across all the condition B. We then test the same 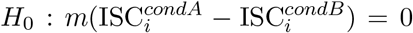. In all cases, the parametric *t*-test are performed after *z*–transformation. Because these tests are performed for each voxel in the brain, we then apply a variety of thresholding methods typically used in fMRI: *p*_FWE_ < 0.05 at the voxel level, *q*_FDR_ < 0.05 at the voxel level, and *p*_unc_ < 0.001, repeating for 1000 random sub samples of *N* subjects. Any subsample with at least one surviving cluster is considered a false positive, yielding an estimate of the false positive rate for that thresholding method with various minimum cluster sizes.

The HBN dataset consists of signals that are actually time-locked due to the presence of a naturalistic stimulus, therefore we needed to employ a slightly different strategy. In this case, for each participant and each movie we extract a window of 100 contiguous volumes, randomly selected within the interval of the full recordings. The choice of 100 volumes is a compromise between having adequately long time series for ISC and being able to properly randomize the segment extraction from limited recordings (250 volumes for TP). To avoid effects due to the beginning and end of the recording we exclude 5 TRs at both extremes of the interval from the selection.

The calculation of leave-one-out ISC consists in computing the Pearson correlation *ρ* between the time course recorded in subject *i* at voxel **v**, with the average time course of the remaining *N* – 1 subjects at the same voxel **v** (Fig. 1a top). We perform this computation with python code based on the package BrainIAK [3]. Note that for the HBN dataset we correlate the activity of one subject watching a random segment of the movie DM with the average brain signal of subjects watching random segments of TP (Fig. 1a bottom). In this way we make sure that no actual synchronization is to be expected in the compared signals, except the fact that they both come as a consequence of processing similar sensory inputs and tasks.

As a result of the ISC computation, we obtain *N* brain maps containing the leave-one-out ISC of each subject. To assess the Type I error rate of parametric tests on this group-level data, we use SPM, one of the most widely used software for statistical parametric mapping [9], to perform second level *t*-tests based on the *r*-to-*z* transformed ISC maps of each participant. For automatizing the analysis for different sample-sizes of subjects and different datasets, we use the python framework provided by nipype [27] in combination with the Gnu parallel command line tool [28]. Importantly, for a given dataset users could perform the same analysis using the SPM GUI, and would only require python code for a single step of the analysis: the ISC computation, which is very simple when using BrainIAK, as explained in [3].

#### 2.2.1. One sample *t*-test

To simulate a situation in which a researcher investigates what brain regions systematically respond to a long stimulus (e.g., a movie) in a given group of participants, we first perform a *t*-test against the null hypothesis *ρ* = 0, separately for all the different group sizes (*N* = 10, …, 50). We *r*-to-*z* transform the ISC maps before applying the *t*-test to ensure normality. We perform this statistical test with and without correction for multiple comparison, and with different minimum cluster sizes. Specifically we used *α_FWE_* < 0.05 corrected at the voxel level and *q_FDR_* < 0.05 corrected at the voxel level. We also present uncorrected data thresholded at *p_unc_* < 0.001. For all of these we test minimum cluster sizes of 1, 5 and 20 voxels. Finally, to assess the effectiveness of the increasingly popular FWE cluster-size correction[29], we also applied that 2 step proceedure, i.e. applying a *p_unc_* < 0.001 cluster-cutting threshold, then extracting the FWEc value, i.e. the minimum cluster size for an FWE cluster-size correction, and then apply that FWEc value on the uncorrected images[29].

#### 2.2.2. Paired sample t-test

To simulate a situation in which a scientist investigates what brain regions respond more systematically to one stimulus than to another, we conducted a matched-pair *t*-test between the *r*-to-*z* transformed ISC calculated in a first segment and the ISC calculated in a non-overlapping second segment of the resting state data of each participant (Fig. 1c). We tested this separately for all the different group sizes (*N* = 10, …, 50). For the HCP datasets, we extract two non-overlapping 200 volume segments from each participant (segment 1 included volumes [100:299], segment 2 volumes [500:699]). For ABIDE, we extracted two 80 volume segments [1:80] and [97:176] from each participant’s fMRI recording. For the naturalistic viewing dataset of the HBN dataset, for each participant we extract a segment of 100 contiguous volumes from a random time window in the movie DM and did the same from the movie TP recordings (Fig. 1c bottom). We then randomly attribute for each participant which of them will be considered condition A and which condition B. Subsequently, we perform the leave-one-out ISC of participant *i* condition A, with the average of condition A of all other participants, then do the same for condition B. After doing so for all participants *i*, we perform a paired *t*-test comparing the leave-one-out ISC for condition A against condition B across the *N* participants.

#### 2.2.3. Two sample t-test

Finally, to simulate a situation in which a scientist investigates what brain regions respond more systematically to a stimulus in one group A of *N* participants (*N* = 10, …, 50) than in another group B of *N* participants (e.g., patients vs. controls) we also performed a two sample *t*-test (Fig. 1b). We randomly selected two non-overlapping subgroups A and B of *N* participants from each dataset, computed the leave-one-out ISC as described for the one-sample *t*-test within each group, and then perform a two-sample *t*-test testing the null hypothesis *ρ*_*A*_ = *ρ*_*B*_ across the two subgroups, again after *r*-to-*z* transforming the data for normality.

### 2.3. Whole brain false-positive quantification

For each type of test, for each of the 1000 random subsamples, and for each combination of sample size, correction type and cluster size threshold, we stored every cluster of voxels surviving the chosen statistical threshold. The proportion of subsamples with at least one significant cluster is then used to estimate the effective Type I error associated with each scenario, and is compared against the 5% to be expected if parametric tests do control adequately for Type I error at the whole brain level. It should be noted that we thus quantify the proportion of analyses that would report any significant ISC cluster anywhere in the brain, as false positive.

### 2.4. True-positive analysis

To assess the sensitivity of the parametric tests, i.e., their ability to detect ISC where there is real correlation, we used the HBN dataset to emulate a situation in which true positives are to be expected.

The strategy here is to define a *ground truth* by performing the leave-one-out ISC on the full sample of 150 subjects separately for the two movies (TP and DM) and compute the threshold with the non-parametric bootstrap method, currently considered optimal for leave-one-out ISC metrics [11]. For each movie we compute the *ground truth* and compare it with the maps obtained from samples consisting of *N* = 10, 20, 30, 40, *or* 50 subjects and thresholded with FWE at *α* = 0.05 and *k* = 5, FDR at *q* = 0.05 and *k* = 5, uncorrected at *p* = 0.001 and *k* = max(20, FWEc) and bootstrap at *p* = 0.001 and *k* = 20. We repeat this operation 50 times for each sample size and movie, computing for each repetition the following metrics for assessing the performance of the tests.

First, we quantify the overall similarity between the thresholded map obtained in each sample and our *ground truth*, estimated with the Cohen *κ* metric.

Second, we determine what proportion of the voxel in the *ground truth* image also survives the thresholding in the sample data (i.e. the hit ratio or sensitivity):

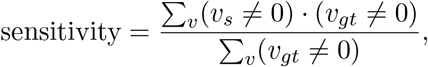

where the sum extends over all voxels *v* and the subscripts *s* and *gt* differentiate thresholded sample and ground truth, respectively.

Finally, we report the proportion of voxels that were not in the thresholded *ground truth*, and that were also not included in the thresholded sample (i.e. correct rejection rate or specificity):

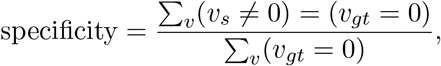

where the sum extends over all voxels *v* and the subscripts *s* and *gt* differentiate sample and ground truth, respectively.

For each of this metric, we combine the results obtained from 50 independent samplings of the movie TP and 50 of the movie DM.

## 3. Results

### 3.1. False-positive analysis

Figure 2 illustrates that for the one-sample *t*-test, i.e. to assess whether ISC is significant in one condition, FDR and FWE corrections for multiple comparisons at the voxel level combined with a minimum cluster size of *k* = 5 voxels or more control false discovery rates to less than 5%. This is true across all the datasets we explored. Inflated Type I errors only occur if no minimum cluster size correction is applied at all, showing that only very small clusters falsely survive this voxelwise threshold. Actually, when applying *k* = 5 as is standard practice in neuroimaging, both FDR (*q* = 0.05) and FWE (*α* = 0.05) corrections are overly conservative, as less than 5% false positives are detected. The same holds true for the stricter requirement on the minimum cluster size *k* = 20, for which no false positives at all are detected.

**Figure 2:**
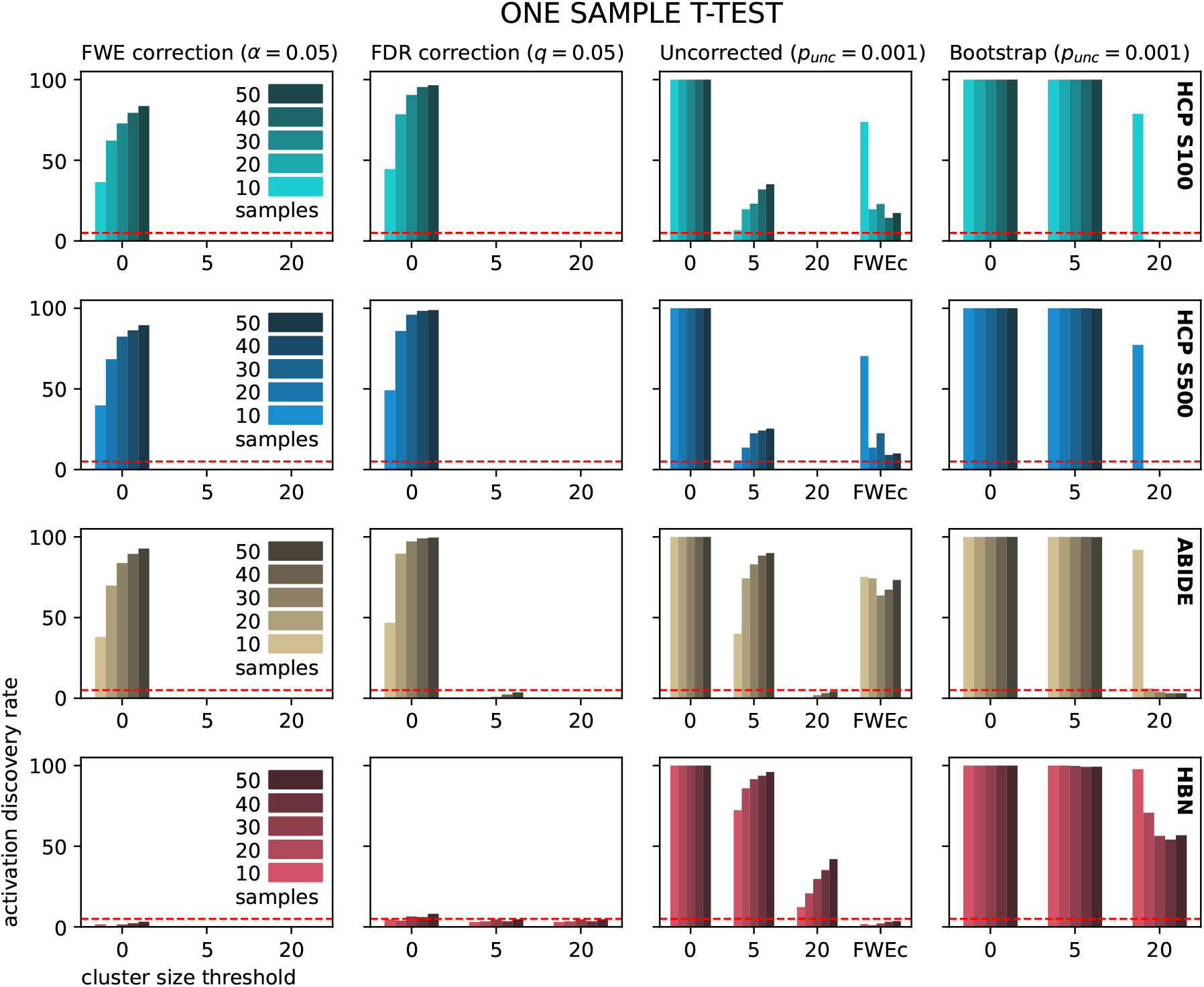
False positives in one-sample *t*-test: percentage of ISC analyses displaying at least one significant cluster when testing against the null condition. In order not to expect any significant cluster, we perform the test on the ISC computed on the brain responses in groups of subjects undergoing resting state (from the top row, HCP S100, HCP S500 and ABIDE) or witnessing naturalistic views in which we randomly mix the data collected at different time points of two different movies (see Sec. 2 for more details). The significance is assessed using (from left to right) voxel-wise Family Wise Error correction (FWE) at *p*_FWE_ < 0.05, False Discovery Rate (FDR) correction at the voxel level at *q*_fdr_ < 0.05, uncorrected *p* threshold at *p_unc_* = 0.001 and non-parametric bootstrap ISC. Every analysis is repeated as a function of minimum cluster size (*x*–axis) and sample size (color bars). For the uncorrected case, FWEc refers to the minimum cluster size that should ensure Family Wise Error correction at the cluster level [29]. The different rows correspond to the datasets HCP S100 (top), HPC S500, ABIDE, and HBN (bottom), respectively. Note that TR = 0.72 s, voxel size = 2 mm for the HCP datasets, TR = 2 s, voxel size = 3 mm for the ABIDE dataset and TR = 0.8 s, voxel size = 2.4 mm for the ABIDE dataset.

Unsurprisingly, applying no correction for multiple comparisons by only applying an uncorrected *p_unc_* < 0.001 threshold is the least conservative of the three analyzed cases. In this case both *k* = 0 and *k* = 5 do not sufficiently control for Type I error over the entire brain. However, imposing stricter thresholds on the minimum clustersize suffices to control Type I error, illustrating again, that only relatively small clusters survive. In particular, for all the resting state datasets *k* = 20 sufficiently controls for Type I errors. Somewhat surprisingly, the FWEc cluster size correction does not reliably limit the false discoveries for this resting state data, and the FWEc estimates obtained with the resting state data were surprisingly small. On the contrary, for the HBN dataset *k* = 20 is still too permissive, whereas the FWEc threshold adequately confines false discovery rates below 5%. The finding that either *k* = 20 or FWEc correction works better in different scenarios might seem counter-intuitive and somewhat arbitrary. However, we can reasonably ascribe this difference to the fact that the four datasets exhibit differences in the spatial smoothness of the imaging data. In particular, we have that the FWHM of the spatial auto-correlation of the ISC data is 1.5 voxels for the HCP datasets, 1.7 voxels for the ABIDE and 3.9 voxels for the HBN, corresponding to 3.0 mm, 5.0 mm and 9.5 mm, respectively. This suggests that for 3T MRI data with a rather small spatial smoothness — less than 2 vox-esl of FWHM in the case of the resting state datasets here examined — it is still advisable to apply a *k* = 20 filter on the cluster size on top of the FWEc correction.

The false positive rates associated with the parametric tests presented in Fig. 2 increase as a function of sample size across the three different datasets. Because the number of participants in all datasets was finite, we wondered whether this increase in false positives as a function of *N* might relate to the fact that as *N* increases, the subsamples will increasingly overlap. However, if this were the case, the effect would be stronger in datasets with fewer participants (HCP S100 and ABIDE) than in the dataset with more participants (HCP S500), but this was not the case. In addition, we performed two simulations where 1000 samples are either drawn from the same data set of 100 simulated brains without systematic correlations, or are drawn independently. Neither of these simulations shows increased false discovery rates with increasing *N* (see Supplementary Material 2).

To assess whether this effect is specific for our parametric tests, we performed the same analyses using a non-parametric bootstrap algorithm in brainiak [11]. Because the issue of multiple comparisons has been less systematically addressed for such non-parametric approaches, here we present the results of the bootstrap method without correction for multiple comparison (Fig. 2), at the same threshold *p_unc_* < 0.001 used for uncorrected parametric analyses. Directly comparing the non-parametric and parametric approaches at the same uncorrected threshold shows that the non-parametric test is slightly less conservative. In particular, with a cluster size threshold of *k* = 5 the fraction of analyses with at least one false positive is always close to 100% in the bootstrap method and consistently higher than in the parametric analyses, particularly for small samples (*N* = 10). When using larger *k* values *k* = 20, for the resting state datasets parametric tests always control Type I error appropriately below 5% while the bootstrap method leads to inflated Type I error rate for small samples (*N* = 10). For the spatially smooth HBN dataset the bootstrap method in combination with the *k* = 20 cluster threshold is not able to control false positives, and currently there are no standard options that allow for applying cluster size corrections to bootstrapped data that would be similar to the FWEc option in SPM.

A salient observation is also that while the parametric tests experience increased false positive rates as *N* increases, the reverse is true for the non-parametric tests, which control Type I errors more effectively as sample size is increased [11]. This highlights the fact that also non-parametric tests such as the bootstrap algorithm used in this case have their weaknesses, particularly when only small samples are available.

Figs. 3 and 4 illutrate the results obtained for scenarios in which two conditions are compared within a given participant and when two different groups of participants are compared, respectively. In these scenarios, we observe that Type I errors are more frequent than for the one sample *t*-test. However, the thresholds that successfully control for Type I errors for the one-sample *t*-tests presented above still control Type I errors quite successfully. For the voxel-wise FWE correction with *α* = 0.05 and *k* = 5, no false positives are ever observed. A FDR correction with *q* = 0.05 in combination with a minimum cluster size of *k* = 5 still controls false positive rates to levels close to the intended 5%. For the paired *t*-tests only the largest sample size *N* = 50 leads to a discovery rate slightly larger than 5% for the connectome datasets. For the two-sample *t*-tests it exceeds the 5% limit only for the ABIDE dataset with a sample size of 10, 20 or 30. When a cluster size threshold of *k* = 20 is used with FDR correction, no false positives are observed at all. In the case of no correction for multiple comparisons, all the observations made for the one sample t-test apply to both the paired and two sample tests. In particular, using a *p_unc_* < 0.001 threshold combined with the larger extent threshold between *k* = 20 or *k* = *FWEc* ensures that the false positive rate remains below 5%.

**Figure 3:**
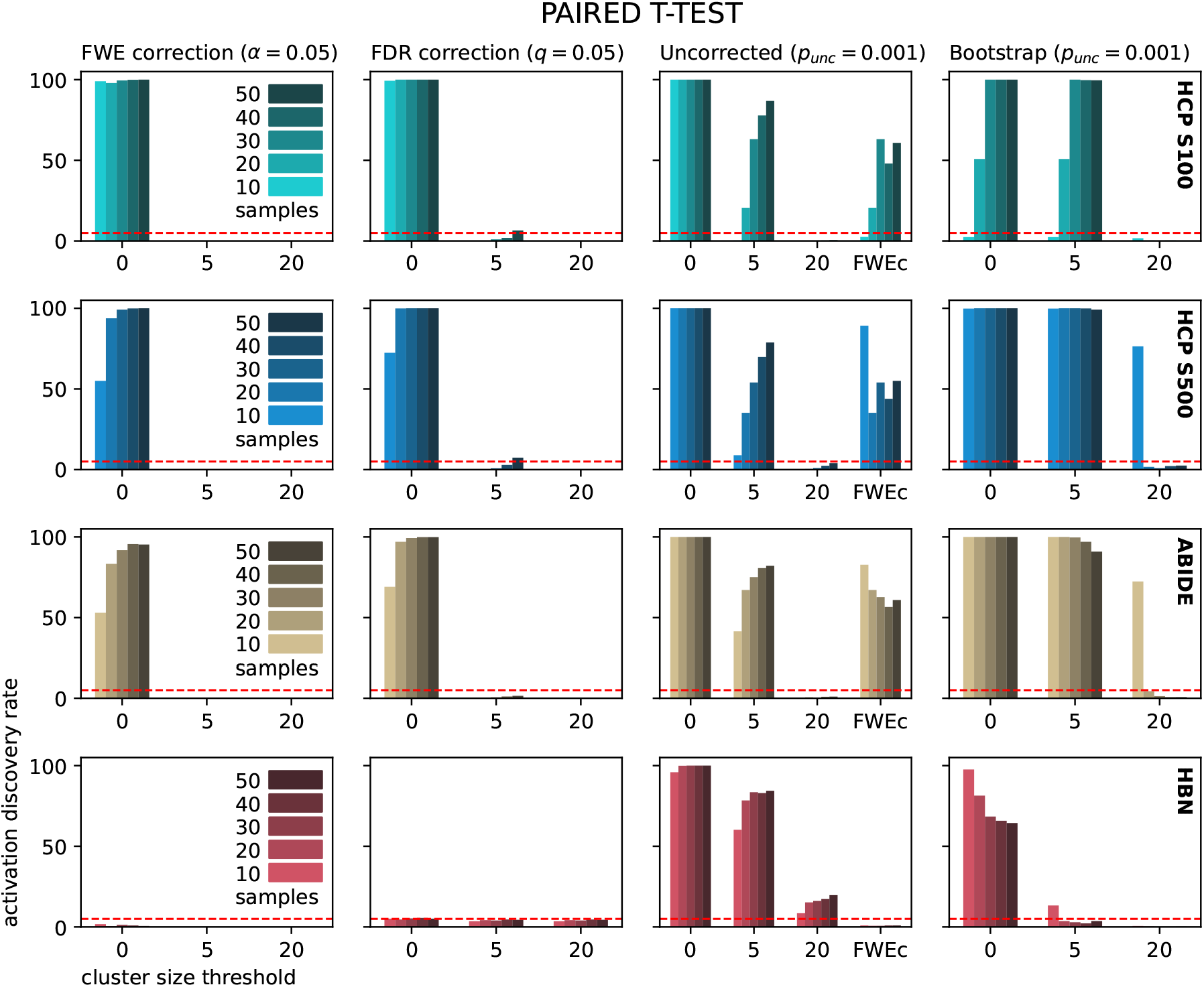
False positives in paired *t*-tests: percentage of ISC analyses displaying at least one significant cluster when testing the difference between pairs of recordings from the same subjects against the null hypothesis. In order not to expect any significant cluster, we perform the test on the ISC computed on the brain responses in groups of subjects undergoing resting state (from the top row, HCP S100, HCP S500 and ABIDE), considering different time windows of the same acquisition as different conditions, or in groups witnessing naturalistic views by comparing the views of different movies as different conditions, randomly flipping the role of each movie (see Sec. 2 for more details). For a detailed description of rows and columns see the caption of Fig. 2.

**Figure 4:**
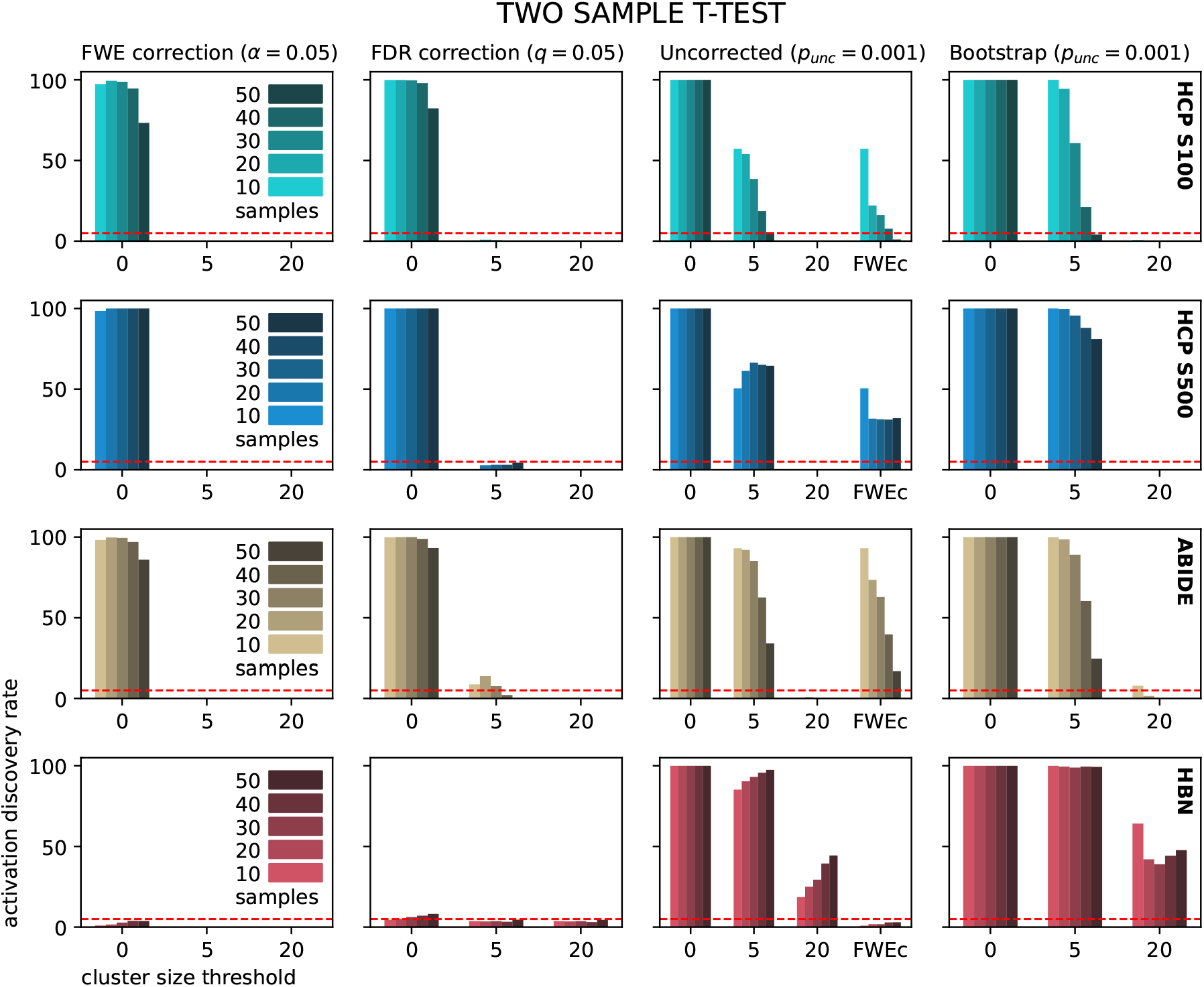
False positives in two-sample *t*-tests: percentage of ISC analyses displaying at least one significant cluster when testing the difference between the population means of two separate groups against the null hypothesis. In order not to expect any significant cluster, we perform the test on the ISC computed on the brain responses in two groups of subjects undergoing resting state (from the top row, HCP S100, HCP S500 and ABIDE) or in two groups witnessing naturalistic views in which we randomly mix the data collected at different time points of two different movies (see Sec. 2 for more details). For a detailed description of rows and columns see the caption of Fig. 2.

Overall, this additional data is consistent with the observations made for the one-sample *t*-test displayed in Fig. 2. However, while false positive rates increased monotonically with sample size for the one-sample *t*-test, this is less clear for the paired *t*-test, and false positive rates actually decrease with increasing sample size in some instances for the two-sample *t*-test (Fig. 4). Regarding the Bootstrap threshold, we observe that this method becomes more effective in the two sample *t*-test, where for *k* = 20 we see a reduced level of false positives compared to the one sample *t*-test, especially noticeable for sample size of 10. This is reasonable as effectively the sample size in a two sample *t*-test is doubled compared to the one sample. We also observe that for the HCP S100 dataset the number of false positives found by the bootstrap method increases with sample size, at least in the range between 10 and 30 subjects, which remains difficult to understand.

### 3.2. True-positive analysis

After verifying that for a one-sample *t*-test FDR and FWE corrections at the voxel level with a minimum cluster size of *k* = 5, as well as *p_unc_* < 0.001 with *k* = *max*(20, *FWEc*) control Type I error rates to below 5%, we explored the relative sensitivity of these thresholding methods to detect ISC in a dataset in which participants did watch the same stimulus. For this purpose we make use of the HBN dataset, which contains imaging data from 150 participants with time-locked brain activity across a wide network of regions involving the processing of the naturalistic view of a movie [19]. We exploit this dataset to compare how different thresholding methods perform in terms of generating results in a subset of the participants that resembles those obtained with the entire dataset using the bootstrap method that is currently considered to generate optimal results[11]. Fig 5 displays Cohen *κ*, sensitivity and specificity of the three selected parametric test and the bootstrap method, as a function of sample size. Note that these values are obtained by pretending that the bootstrap method applied to the entire dataset of 150 subjects (’Bootstrap150’) identifies the ’true’ network of brain regions that time-lock to the stimulus. However, given that the bootstrap method is unlikely to have perfect sensitivity, some voxels of the true network will not be in the Bootstrap150 map. If in such a voxel, one of the parametric methods comes to a significant result, this should contribute to increasing the measured sensitivity of the parametric method but will, instead, erroneously reduce the measured specificity of the method. Hence, our measures should be interpreted with this caveat in mind. As one could expect, Cohen *κ* and sensitivity increase with sample size, whereas specificity decreases. Averaging over the 50 realization of different samples performed per each movie, the FDR method with *q* = 0.05 and *k* = 5 scores the highest values for both Cohen *κ* and sensitivity, for all sample sizes. However, this is the method that shows the least specificity, with values as low as 80% for *N* = 50 participant samples. A threshold with FWE correction at *α* = 0.05 and *k* = 5 is the most conservative, with a sensitivity that reaches only 20% for a sample size of 50 and a specificity that is nearly always 100%. Uncorrected *t*-test with *p_unc_* = 0.001 and *k* = max(20, FWEc) and bootstrap with *p* = 0.001 and *k* = 20 fall somewhere in between the aforementioned methods, with the bootstrap being slightly more conservative.

**Figure 5:**
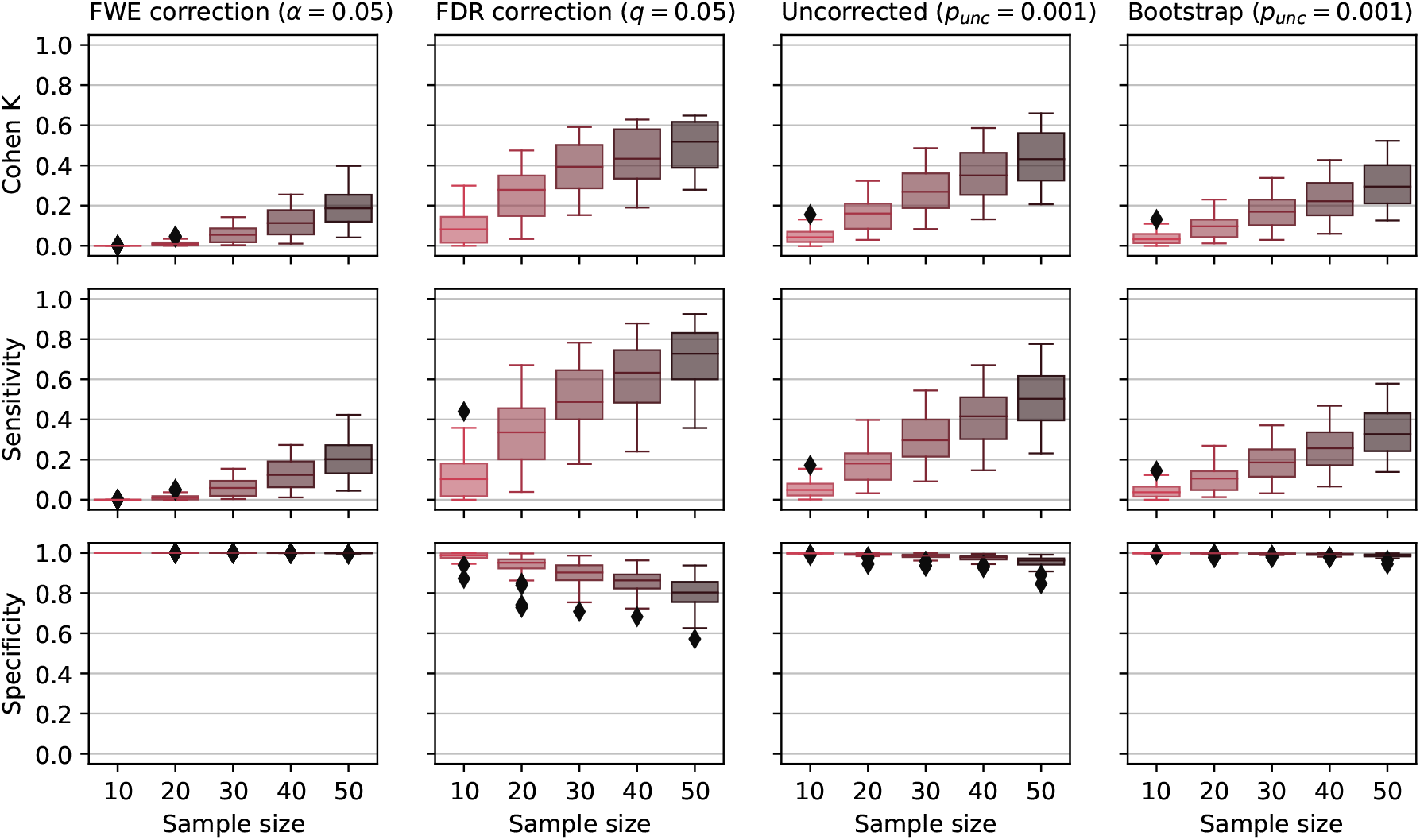
Overview of sensitivity and specificity of the four selected methods for thresholding ISC data. From left to right, FWE correction with *α* = 0.05 and *k* = 5, FDR correction with *q* = 0.05 and *k* = 5, uncorrected threshold with *p_unc_* = 0.001 and *k* = max(20, FWEc), bootstrap threshold with *p* = 0.001 and *k* = 20. The three rows correspond from top to bootom to Cohen *κ*, sensitivity and specificity. The boxplot summarizes statistics from 50 independently drawn samples for each movie (100 samples in total).

Although the FDR method is the most sensitive and the one scoring the best values of Cohen *κ*, its data for specificity are at first sight worrying. However, as noted above, our ability to produce a ground truth data is also limited, as we are assuming the bootstrap method with 150 subjects *p* = 0.001 and *k* = 20 to be an ideal scenario, which does not need to be the case.

Comfortingly, when looking at the maps produced by the different methods, we observe that most of the “false positives” elicited by the FDR method are located in the same clusters as the “true positives”. This can be seen in Fig. 6, where we display a representative horizontal slice with the significant voxels determined by each thresholding method in one computation of ISC in the TP movie, for a sample size of 20 (above) and 50 (below). The heatmaps represent *t*-values for the parametric methods and median r for the bootstrap. These are overlaid on a green map that represent the Bootstrap150 ground truth data. By eye, the results of the different thresholding methods all agree in locations where ISC is highest (lighter colors), but disagree in the extent of the networks that are detected as synchronizing their activity to the stimulus. We can clearly see that the FDR threshold is the most sensitive among the four algorithms, and examining the top row (*N* = 20) shows that FDR is the only method sensitive enough at the sample size of typical fMRI studies to detect the prefrontal synchronization that can otherwise only be detected in large sample sizes. At the same time, FDR does not yield to significant ISC in regions far from those included in the Bootstrap150 *ground truth*.

**Figure 6:**
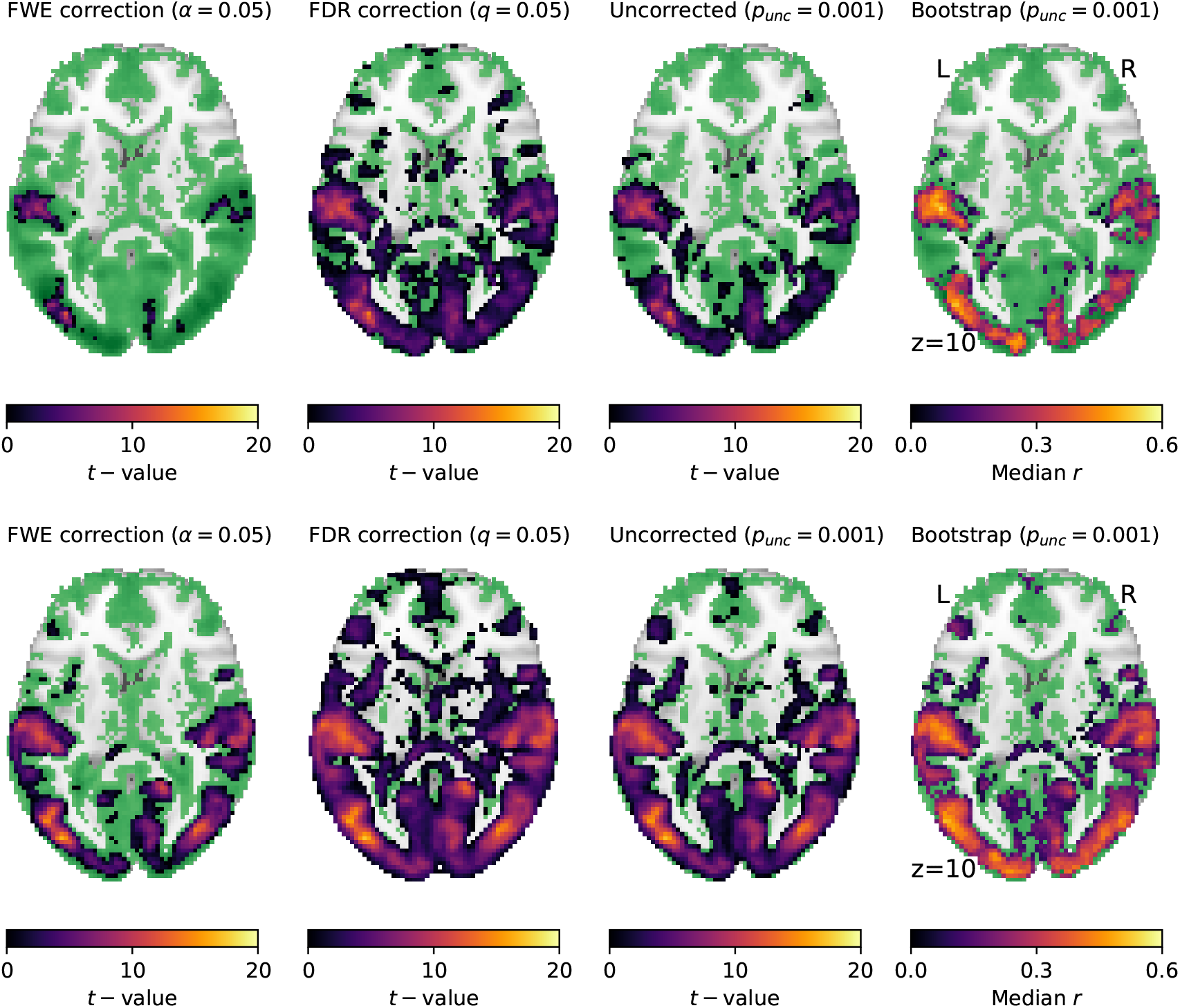
Brain maps for the true positive analysis. Example of a horizontal slice from one repetition for each of the investigated threshold method (warm-colored heatmap), compared to the *ground thruth* (Bootstrap150 in green), for *N* = 20 (above) and *N* = 50 participant subsamples (below). For each method we show significant voxels with corresponding *t*-value (for the parametric) or median r (for the bootstrap) as heatmaps, directly overlayed on the Bootstrap with 150 samples (light green). The images show a representative slice located in MNI coordinates at *z* = 10.

## 4. Discussion

In this work we evaluate the performance of parametric *t*-tests in the context of leave-one-out Inter-Subject Correlation based on publicly available resting state and task based datasets. We find that parametric tests is a valuable alternative to non-parametric testing by offering an appropriate protection against false positives in the absence of true ISC and superior sensitivity in the presence of true ISC.

To arrive at that conclusions, we first explored whether parametric tests appropriately control Type I error, i.e., how often they would falsely detect significant synchronization in resting state datasets or in a task based dataset in which the timing and content of the presented naturalistic stimulus is unmatched. Our results indicate that parametric tests control Type I errors when testing hypotheses on ISC data in ways that are quite similar to the way they control Type I errors in standard subtraction designs. In particular, FDR corrections (*q_FDR_* < 0.05) at the voxel level paired with a minimum cluster size of *k* = 20 voxels robustly protected against the false detection of synchrony in all the resting state datasets and types of comparisons we explored. This was true despite the actual differences in voxel-size and smoothing used in the HBN, HCP and ABIDE. Because resting state data may have signal properties that do not approximate those of the naturalistic viewing conditions in which ISC is typically used, it is important that FDR with k=20 also protected against excessive false-positive rates in the face of data that comes from naturalistic viewing, but where the temporal locking is disturbed. Importantly, this thresholding method not only protected against false positives: it was also the most sensitive in detecting synchrony in datasets in which participants watch a common stimulus, and indeed, was the only method sensitive enough to correctly detect synchronization in prefrontal cortices even in the sample sizes typically used in cognitive neuroscience studies.

FWE corrections at the voxel level (*α_FWE_* < 0.05) also controls Type I error rates very effectively, even when a minimum cluster size of only *k* = 5 voxels is used. However, FWE correction was the least sensitive of the methods we explored, when detecting synchrony in participants viewing the same movie. FDR correction thus appears to offer the best balance between Type I and Type II error, and may be preferable when scientists investigate smaller effect sizes in modest sample sizes - a situation that is unfortunately typical of most cognitive neuroscience studies.

Using uncorrected (*p_unc_* < 0.001) thresholds in combination with a cluster size that is defined as the maximum between *k* = 20 and *k* = *FWEc* also controls Type I error while providing reasonable sensitivity. Surprisingly, simply using *k* = *FWEc*, without a minimum of k=20, did not always work well. In particular, for the resting state data we explored, where smoothness is very low, FWEc estimates were insuficiently small.

Our study has a number of limitations. Firstly, finding that a method controls against Type I error in a particular dataset does not ensure that it controls against Type I error in all datasets. We therefore challenged our parametric thresholding methods with 4 different datasets, which varied in TR values (ranging from 0.72s to 2s), isotropic voxelsize (ranging from 2 mm to 3 mm), smoothness (ranging from 3 mm to 9 mm), and acquisition schemes (multiband acceleration in the HCP and HBN but not the ABIDE). Importantly, we also exposed our candidate methods to data that comes from resting state and naturalistic viewing tasks. Finally, we varied our (resampled) sample sizes from 10 to 50 participants. That our parametric thresholding methods performed well across all these datasets suggests that parametric tests robustly control for Type I error across a reasonable range of parameters. Additionally, we also explored whether our results depend on the length of the neuroimaging data we consider, but our results are robust over a wide range of segment length (see Supplementary Material 1). Secondly, the assumption that neuroimaging data does not include any synchronization between participants might be doubtful, even for the case of resting state acquisitions. Because data was taken at the same moment relative to the start of data acquisition in all participants, some synchronization may occur across participants in reaction to sensory and affective processes triggered by being scanned in the same fMRI scanner. That parametric tests control against finding synchronization across participants despite this possibility is reassuring. Thirdly, that the additional voxels detected by FDR thresholding fall outside of the Bootstrap150 results (that we use as a proxy of the ground truth) does not guarantee, that these voxels are truly false positives. Accordingly, the specificity drop we observe in large samples with FDR may actually represent increases in sensitivity, but we are unable to assess that.

Importantly, our results only apply to leave-one-out ISC. Naturally, parametric tests cannot be straightforwardly used on pairwise ISC data, where the numbers of degrees of freedom corresponding to the number of measured subjects is dramatically different from the number of pairs that one can generate with such subjects.

It is our experience that many neuroimagers stay away from using ISC-based neuroimaging paradigms because they shy away from the relatively complex coding necessary to perform robust statistical inference using non-parametric tools. We trust that our results will allow a broad community of neuroimagers trained in using and interpreting parametric second level neuroimaging statistics implemented in SPM or FSL to leverage the conceptual power of ISC. This is because when using parametric statistics, ISC data can be analysed using a pipeline almost identical to that of traditional fMRI analysis, including standard preprocessing, and standard second level analyses. Only the actual calculation of the ISC needs to be done using less standard code, but this step can be easily implemented using toolboxes we provide in [3]. Our findings also naturally extend to more sophisticated forms of ISC, including inter-subject functional connectivity, that rely on very similar, leave-one-out computations [3]. In addition to making analyses easier to run, the use of parametric statistics implemented in standard packages in particular paves the way to the systematic usage of sensitive but robust corrections for multiple comparisons (i.e., FWE and FDR) that adapt to the number of voxels and the spatial smoothness of a dataset. Such adaptive corrections are not currently readily available for non-parametric approaches. Finally, non-parametric approaches for ISC can become prohibitively demanding in terms of computations as sample sizes increase, and parametric tests can thus further improve the efficiency of data analyses. While we by no means wish to argue against non-parametric testing of ISC data, we hope that our validation of parametric tests will provide scientist with an easier-to-implement yet powerful and sensitive method to complement existing non-parametric tests.

## Acknowledgments

We thank Christiaan Becker and Pierre-Louis Bazin for performing a pilot analyses that preceeded and informed this study, Ritu Bhandari for insights on the use of SPM, Alessandra Nostro and Leonardo Cerliani for help to identify appropriate resting state datasets.

