## Supplementary Material for "Parametric tests for Leave-One-Out Inter-Subject Correlations in fMRI provide adequate Type I error control while providing high sensitivity"

### 1. False positives as a function of temporal window

Our choice of using 200 TRs from the HCP datasets was somewhat arbitrary. To test the impact this decision has on our Type I error, we repeated our one-sample t-test analysis for a fixed group size of $N = 20$ but varying the number of volumes included from 10 to 400 TRs. Figure 1 shows that FWE ($p_{FWE} < 0.05$, $k = 0.05$) and FDR ($q = 0.05$, $k = 5$) protect against Type I error regardless of the number of volumes included. When detecting nonzero false positive rates (using no minimum cluster size), we notice an increase in Type I error up to approximately 100 TRs, followed by a decrease as larger segments are considered. These results are also important for applications in which short temporal windows are necessary, because analyses attempt to identify moments within a longer stimulus in which a particular network is selectively engaged.

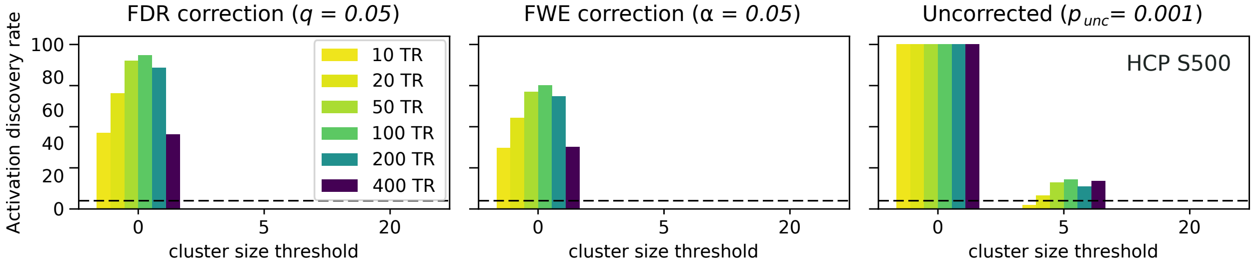


Figure 1: Analysis of the false-positive rate for one-sample t-test on intersubject correlation data at varying the temporal window used for the computation of the leave-one-out isc.

* Corresponding author
** These authors contributed equally to this work

### 2. Simulations for estimating the effects of sampling from a finite dataset

To check whether sampling 1000 times with different sample size from a finite dataset can lead to artifacts in the false positive rate measured in the main text, we performed numerical simulations. We created two scenarios. In the first scenario, similar to our experimental framework, we generate a dataset of 100 simulated brain data and compute the ISC by sampling this data 1000 times per each sample size. In the second scenario, we compute ISC starting from simulated data which is independently generated at each of the 1000 iterations. To mimic resting state data, we generate simulated data as Gaussian random noise with a finite temporal and spatial correlation. From Fig. 2 we deduce that having a limited dataset consisting of 100 independent measurements does not produce visible effects on the false positive analysis, as the results for the two different scenarios (top and bottom panels, respectively) are consistent with each other.

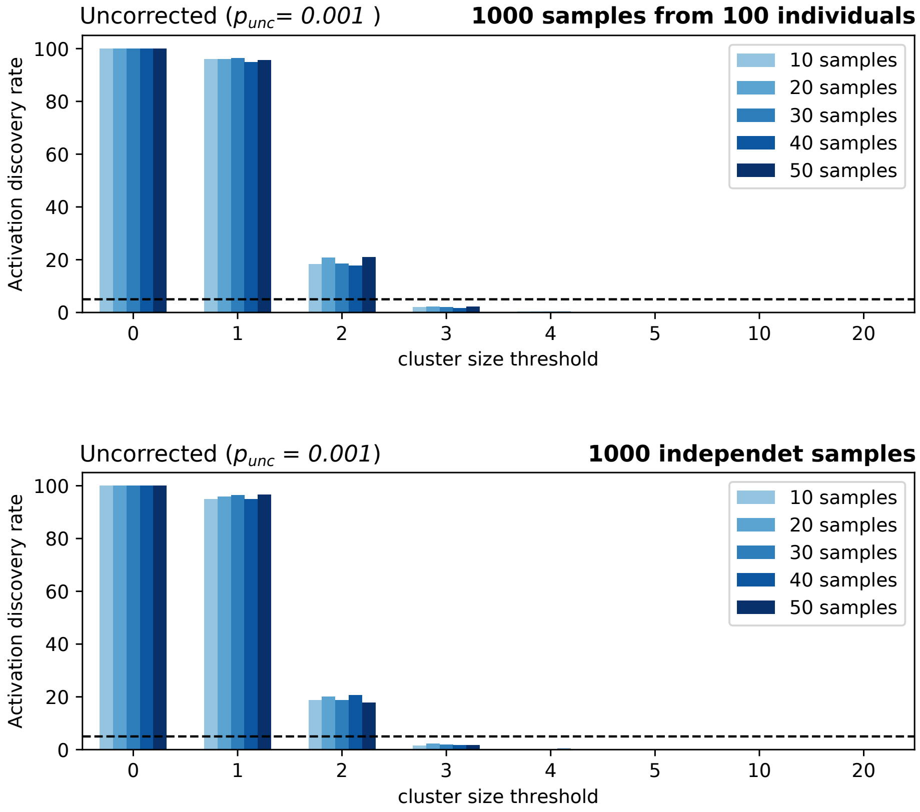


Figure 2: One sample t-test for simulated brain activity. We compute the ISC starting from simulated activity consisting in gaussian random noise with a finite temporal and spatial correlation. In the top panel a scenario similar to the experimental one, in which only 100 brains are available for random sampling. In the bottom panel a scenario in which the brain activities are independently generated for every step contributing to the overall statistics.
